# An Assumption of The Regulatory Function of Nf2-Amot Complex in Early Mammalian Embryos with A Computational Model

**DOI:** 10.1101/2024.03.31.587462

**Authors:** Yusuke Sakai, Jun Hakura

## Abstract

The paper assumed that Nf2-Amot complex regulates the phosphorylation cascade so that each cell in the early mammalian embryo differentiates properly *in silico*. To confirm the validity of the assumption, it was necessary to verify whether Nf2-Amot complex has an impact on the resulting differentiation. The living embryo is unsuitable for the confirmation since the early mammalian embryo is too small to observe and too ethically sensitive to invade. In such cases, computational models can be used as experimental subjects for operations that cannot be applied to the living embryo. Previous models on the embryo, however, could not verify the assumption because they had not modeled Nf2-Amot complex, and they seldom modeled the Hippo signaling pathway. Therefore, the paper introduced a model of Nf2-Amot complex to the previous study that had modeled the Hippo signaling pathway. Testing the model under diverse conditions revealed that the existence of Nf2-Amot complex reproduces the ideal cell differentiation observed in the living embryo. In this sense, the validity of the model was confirmed. Furthermore, diverse cell-cell contacts that induce various concentrations of Nf2-Amot complex also resulted in ideal cell differentiation. These results suggested the correctness of the assumption *in silico*.

## INTRODUCTION

Cell differentiation is a carefully orchestrated process in which cells form to fulfill their roles according to their dynamic location in an organism. Especially, the early mammalian embryo is a suitable system for studying because it differentiates into just two simple states, i.e., trophectoderm (TE) or inner cell mass (ICM). These mechanisms are expected to be able to apply to complex structures in the following developmental stages of the embryo.

During the early developmental process, each cell in the embryo recognizes its position based on the states of cell-cell contact. The first differentiation event occurs from the 8-cell stage. The process is caused by the existence of the Par-aPKC complex (Par-3, Par-6, atypical protein kinase C) and E-cadherin [Ohno, 2001; Suzuki et al., 2006; Rubtsova et al., 2022; Vries et al., 2004]. These proteins induce Cdx2 (caudal type homeobox 2) and Oct4 (octamer-binding transcription factor 4) productions in the embryo [Hirate et al., 2015; Redmer et al., 2011; Sasaki, 2015]. While every cell-cell contact surface produces E-cadherin, the noncontact surface of a cell produces Par-aPKC. The concentrations of Cdx2 and Oct4 play important roles in determining whether cells differentiate into trophectoderm (TE) or inner cell mass (ICM) [Strumpf et al., 2005]. Namely, the supremacy in the promotion of Cdx2 expression results in TE, and the promotion of Oct4 expression results in ICM [Sasaki, 2017]. The expression of Oct4 and Cdx2 are the results of activation/inactivation of the Hippo signaling pathway, respectively (**Fig.1**). The Hippo signaling pathway is the phosphorylation cascade that activates Amot (angiomotin), Nf2 (Neurofibromin type 2), Lats (protein kinase Lats) and Yap (yes-associated protein), step by step [Pan, 2010; Zhao et al., 2007; Hirate et al., 2014; Wu et al., 2021]. In the outer cells, Amot is trapped by Par-aPKC organized in an apical domain [Hirate et al., 2013; Ohno, 2001]. This inhibits the activation of Lats, then allowing Yap to enter the nucleus. In the nucleus, the bond of Tead4 (the TEAD/TEF family transcription factor Tead4) and Yap promotes Cdx2 expression and inhibits Oct4. On the contrary, phosphorylated Amot and Nf2 form a Nf2-Amot complex on E-cadherin in the inner cell. The Nf2-Amot complex promotes phosphorylation of Lats [Yu et al., 2013; Cockburn et al., 2013; Yin et al., 2013; Li et al., 2015; Saiz et al., 2020]. The phosphorylation of Lats also causes phosphorylation of Yap, cytoplasmic localization, and degradation of Yap [Kim et al., 2011]. In the nucleus, the expression of Cdx2 is downregulated, and the expression of Oct4 is upregulated [Toyooka, 2020; Nishioka et al., 2009].

**Fig. 1.**
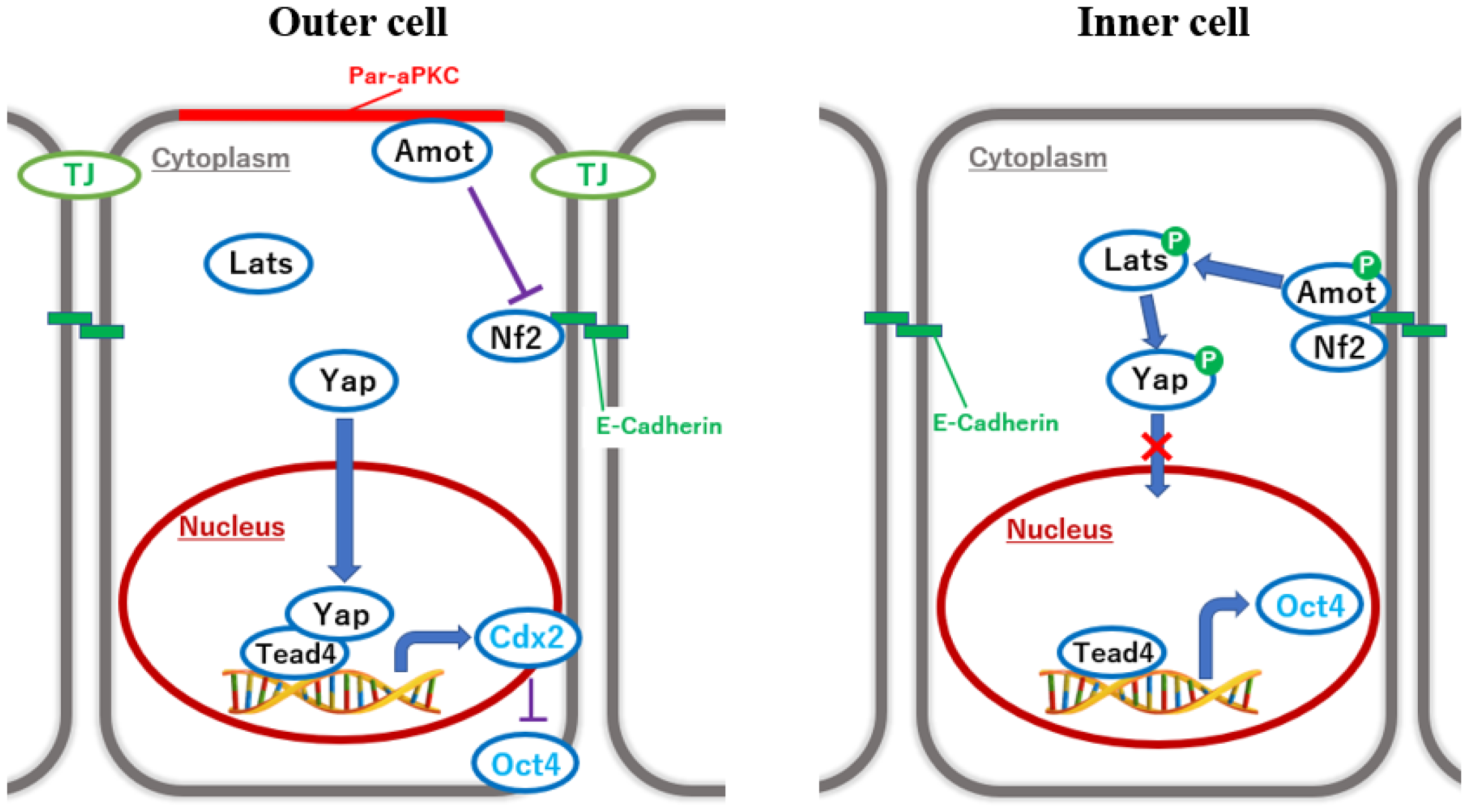
Differentiation mechanisms in the early mammalian embryo. The Hippo signaling mechanisms regulate the separation of TE and ICM. The outer cell differentiates into TE (left). The cell does not activate the Hippo pathway, thus allowing Yap to enter the nucleus and promote Cdx2 expression. The inner cell differentiates into ICM (right). The cell activates the Hippo pathway, phosphorylates, degrades Yap, and inhibits Cdx2 expression.

The embryo *in vivo* almost always succeeds in its differentiation even if its cells happen to change their positions, caused by cell divisions and cell migrations. The paper defines the term robustness as the adapting ability of the embryo to the diversity of cell-cell contacts in the embryo. During the formation of the TE/ICM, cells divide and migrate [Watanabe et al., 2014; Yamanaka et al., 2010], and seldom remain at their original position in the embryo. In this sense, cell-cell contact conditions dynamically change, and the cell differentiation mechanism is necessary to be robust to adapt to the conditions mainly related to the position of each cell [Cang et al., 2021]. This robustness is achieved by the regulation of phosphorylation. [Bae et al., 2018]. Nevertheless, few details of the mechanism are revealed, since the size and inaccessibility of the embryo [Shahbazi et al., 2019]. Therefore, studies on cell differentiation often use theoretical approaches [Herrera-Delgado et al., 2021; Cockerell et al., 2023; Geard et al., 2009] by introducing a model to examine new insights.

Among computational models, the previous model proposed by Caluwé et al focuses on the Hippo signaling pathway and conducts modeling [Caluwé et al., 2019]. The previous model focused on how the Hippo signaling controls the Cdx2/Oct4 expression. The model expresses the positions of cells in the embryo by concentration dynamics of Par-aPKC and E-cadherin. Since the both status and timing of each cell-cell contact are different, the concentration dynamics of Par-aPKC and E-cadherin should be different. However, the paper by Caluwé et al showed only one temporal evolution of the concentration dynamics for the outer cell and the inner cell, respectively. We, therefore, conducted a preliminary experiment assuming the diverse conditions on cell-cell contacts. The results showed that position-specific cell differentiation is expressed only in specific cell-cell contact conditions.

Gene Regulatory Network (GRN) modeling is a well-established theoretical approach that has been used to infer or understand many developmental mechanisms. The GRN models configure the interactions between the specific transcription factors. In the GRN models, the concentration dynamics are represented by temporal evolutions of protein concentrations and toward one of the multiple stable steady states [Robert et al., 2022; Wu et al., 2013; Mot et al., 2016]. Thus, GRN models are frequently used to infer the role of target substances, e.g., genes or proteins, in the field. Most studies on early development use differential equations with GRN models, to allow the investigation and analysis of continuous time courses [Chickarmane et al., 2006; Holmes et al., 2017].

The paper focuses on the role of the Nf2-Amot complex and quantifies its effect on robustness. To make the difference between the previous and the proposed models clear, we represent them as forms of GRN models as in **Fig.2**. In the previous model, the gene regulatory network of Cdx2/Oct4, consists of the following eight components: Par-aPKC, E-cadherin, AmotM (Amot trapped by Par-aPKC), AmotP (phosphorylated Amot), LatsP (phosphorylated Lats), Yap, Cdx2, and Oct4 (**Fig.2A**). However, it does not consider the Nf2-Amot complex, which regulates the phosphorylation cascade in the Hippo signaling pathway [Hong et al., 2020; Shi et al., 2017; Lanner, 2014]. Thus, the proposed model integrates the Nf2-Amot complex into the previous model (**Fig.2B**) to make cells more robust to their position change and succeed in differentiation. To achieve this, we introduce the dynamics of the Nf2-Amot complex described by the differential equations based on the previous model proposed by Caluwé et al. Then, we compared these models to examine the role of the Nf2-Amot complex. The results show how the Nf2-Amot complex effectively works, more to say, indispensable to achieve robustness in the differentiation.

**Fig. 2.**
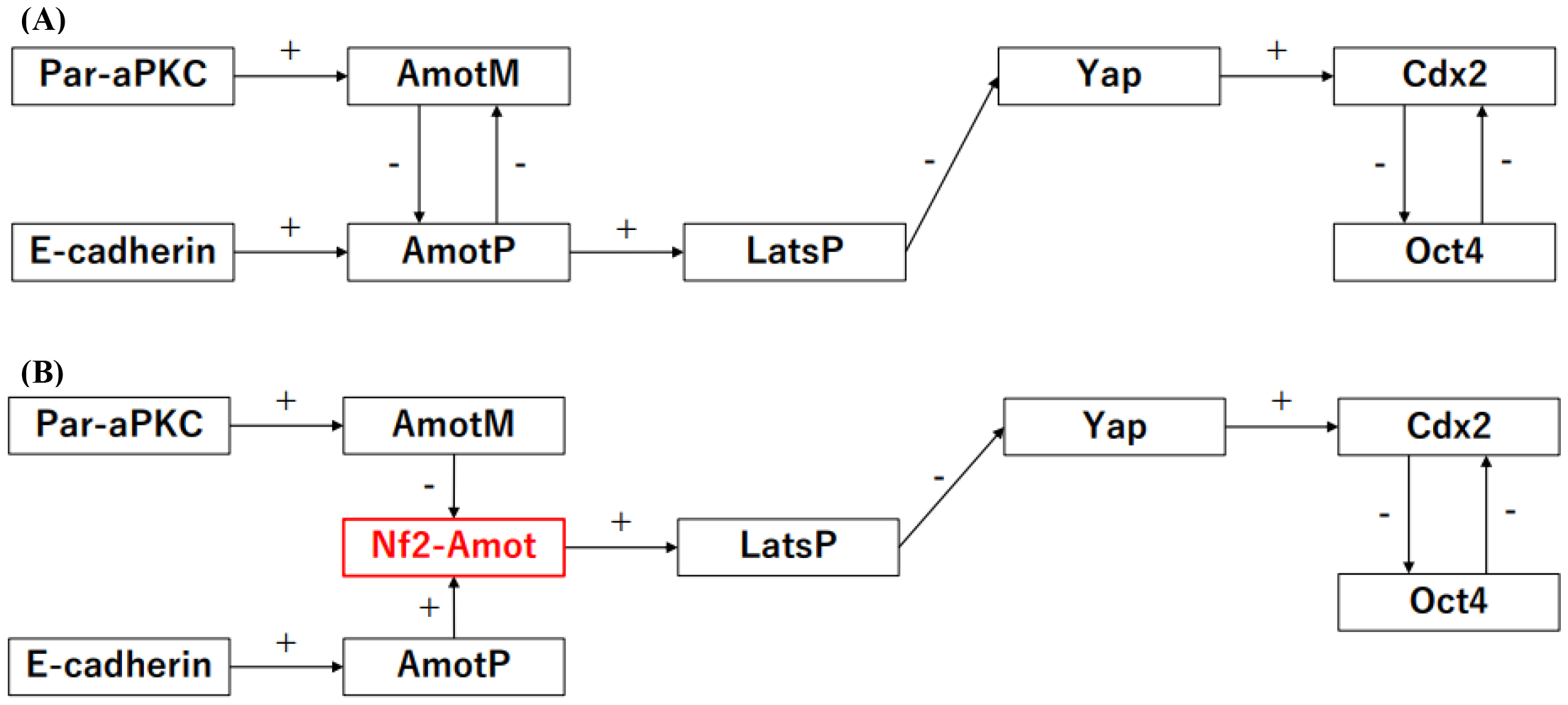
Gene regulatory network models of intracellular mechanisms. The figure depicts the difference among the components in the computational models. Where “+” reflects promotion, and “-” reflects inhibition. (A) The previous model: AmotP (Phosphorylation of Amot) promotes LatsP (Phosphorylation of Lats). (B) A proposed model: the Nf2-Amot complex promotes LatsP.

The main purpose of the study was to understand the key regulatory function of phosphorylation to improve robustness. The paper, therefore, emphasizes computational models. On the other hand, from the biological perspective, the paper comes to reach a novel biological assumption that the Nf2-Amot complex regulates the phosphorylation cascade of the Hippo signaling pathway to achieve robust cell differentiation in early mammalian development.

## RESULTS

To investigate the regulatory mechanism of the Hippo signaling pathway, we construct a model of the phosphorylation cascade in early mammalian embryos. We, first, describe the proposed model using ordinary differential equations (ODEs) and a parameters selection in the model of the Nf2-Amot complex. The proposed model integrates the Nf2-Amot complex into the previous model to improve robustness. In the proposed model, Equations (1) to (4), (7), and (8) are the same as the previous model. The proposed model adds Equation (6) which represents the dynamics of the Nf2-Amot complex. In Equation (6), the ranges of parameters are determined empirically by considering the previous model. Within the ranges, we select one combination of parameters through an experiment described in the later subsection. Equation (5) expresses phosphorylated Lats influenced by the Nf2-Amot complex. Furthermore, the paper proposes concentration dynamics of Par-aPKC/E-cadherin (Equations (9) and (10)) to simulate the diverse cell-cell contacts. These proposed models use the dynamics of the previous model as references. We, then, conduct two experiments: inhibition of the Nf2-Amot complex, and comparison of the proposed model and the previous model. The former experiment confirms whether the dynamics of the proposed model follow that of the embryo *in vivo*. The latter experiment examines how the existence of the Nf2-Amot complex influences the expressions of Cdx2 and Oct4.

### Computational modeling of cell differentiation during early mammalian development

TE/ICM differentiation can be inferred from the amount of Cdx2 and Oct4 expression. Cdx2 and Oct4 mutually inhibit each other, and eventually one of the proteins is highly expressed. In the computational models, Caluwé et al have proposed ODEs to describe the Cdx2/Oct4 transcriptional system based on the observation of a phosphorylation cascade in the Hippo signaling pathway. The previous study has determined TE/ICM differentiation depending on which protein has a higher level during the multiple stable steady states. Equations (1) and (2) express the concentration dynamics of Cdx2 and Oct4 in cell *i*, respectively [Caluwé et al., 2019]. In the first term, while the first bracket represents self-amplification, the second bracket represents mutual inhibition. The second term represents protein degradation:

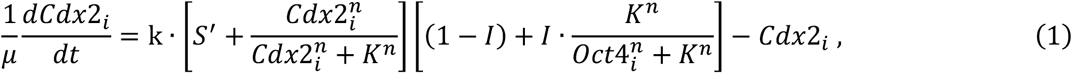

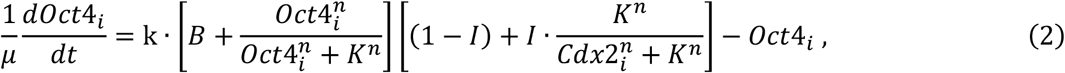

where the variable *Cdx*2_*i*_ represents an expression level of Cdx2 in cell *i*, and the variable *Oct*4_*i*_ represents an expression level of Oct4 in cell *i. i* represents an identifier of a cell. Furthermore, *n* represents the Hill coefficient of auto-activation and cross-inhibition. The Hill coefficient is the measure of how steep the response curve is. While parameter *μ* represents the timescale, we adjust the parameter *μ* to introduce the model for the Nf2-Amot complex afterward. The other parameters are explained in Table 1.

**Table 1.**
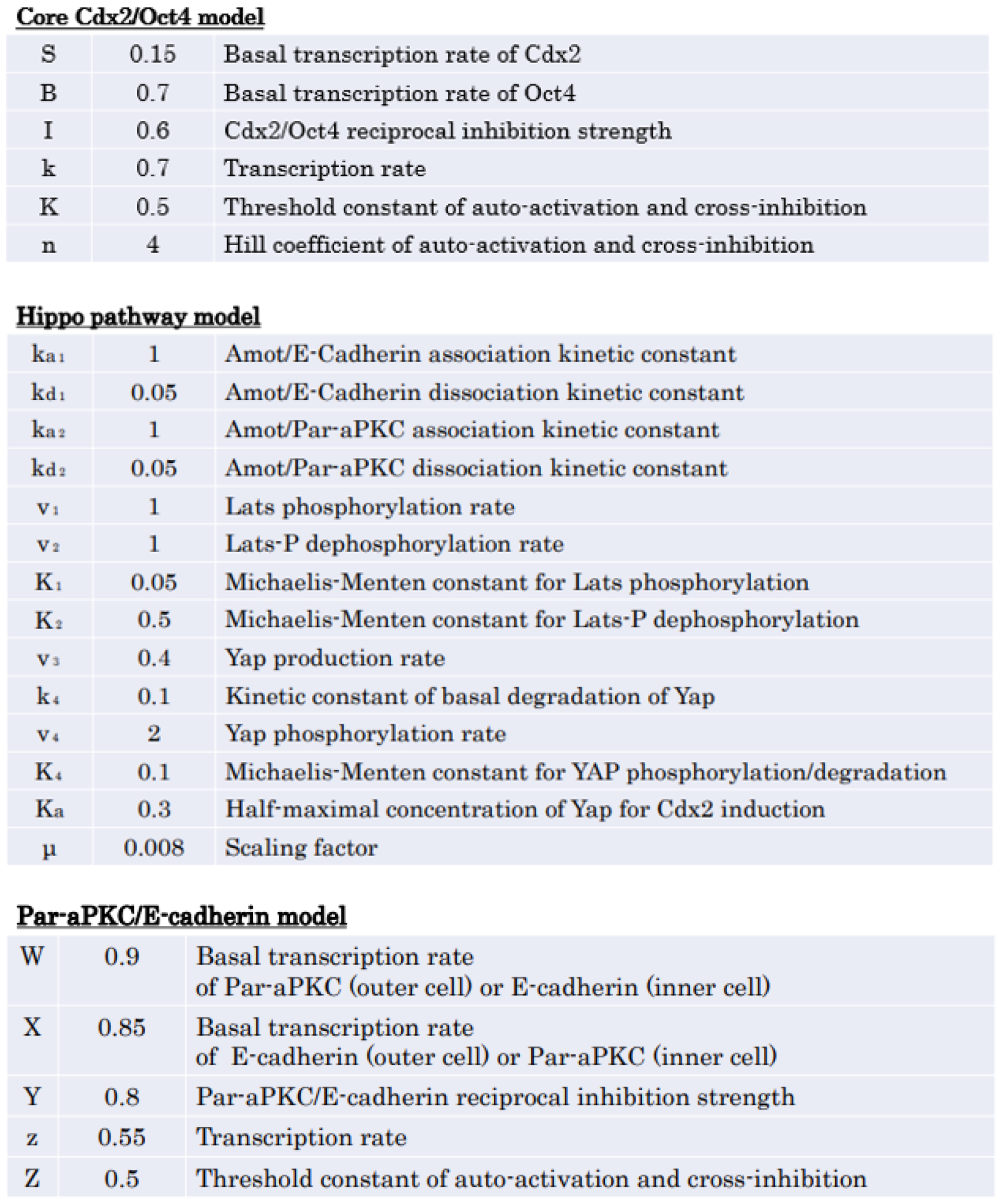
List of parameters in the proposed model. The parameters of the Core Cdx2/Oct4 model and the Hippo pathway model are the same as the model of the previous study [Caluwé et al, 2019], except for μ. We adjust the parameter μ to introduce the model for the Nf2-Amot complex. Furthermore, the paper proposes the parameters of the Par-aPKC/E-cadherin model. These parameters are based on dynamics of Par-aPKC/E-cadherin which is represented by temporal evolutions in the previous study.

**Core Cdx2/Oct4 model**
|  |  |  |
| --- | --- | --- |
| S | 0.15 | Basal transcription rate of Cdx2 |
| B | 0.7 | Basal transcription rate of Oct4 |
| I | 0.6 | Cdx2/Oct4 reciprocal inhibition strength |
| k | 0.7 | Transcription rate |
| K | 0.5 | Threshold constant of auto-activation and cross-inhibition |
| n | 4 | Hill coefficient of auto-activation and cross-inhibition |
| **Hippo pathway model** |  |  |
| $k_{a1}$ | 1 | Amot/E-Cadherin association kinetic constant |
| $k_{d1}$ | 0.05 | Amot/E-Cadherin dissociation kinetic constant |
| $k_{a2}$ | 1 | Amot/Par-aPKC association kinetic constant |
| $k_{d2}$ | 0.05 | Amot/Par-aPKC dissociation kinetic constant |
| $v_1$ | 1 | Lats phosphorylation rate |
| $v_2$ | 1 | Lats-P dephosphorylation rate |
| $K_1$ | 0.05 | Michaelis-Menten constant for Lats phosphorylation |
| $K_2$ | 0.5 | Michaelis-Menten constant for Lats-P dephosphorylation |
| $v_3$ | 0.4 | Yap production rate |
| $k_4$ | 0.1 | Kinetic constant of basal degradation of Yap |
| $v_4$ | 2 | Yap phosphorylation rate |
| $K_4$ | 0.1 | Michaelis-Menten constant for YAP phosphorylation/degradation |
| $K_a$ | 0.3 | Half-maximal concentration of Yap for Cdx2 induction |
| $\mu$ | 0.008 | Scaling factor |
| **Par-aPKC/E-cadherin model** |  |  |
| W | 0.9 | Basal transcription rate of Par-aPKC (outer cell) or E-cadherin (inner cell) |
| X | 0.85 | Basal transcription rate of E-cadherin (outer cell) or Par-aPKC (inner cell) |
| Y | 0.8 | Par-aPKC/E-cadherin reciprocal inhibition strength |
| z | 0.55 | Transcription rate |
| Z | 0.5 | Threshold constant of auto-activation and cross-inhibition |

The expression of Cdx2 requires Yap to enter the nucleus, and then Yap binds Tead4 in the nucleus. The model does not consider Tead4 to focus only on the influences of Yap. We, therefore, intend to ignore additional substances other than the Nf2-Amot complex. To reflect Yap in the production of Cdx2, Equation (3) expresses basal rates *S*^′^ in the expression of Cdx2 as [Caluwé et al., 2019]:

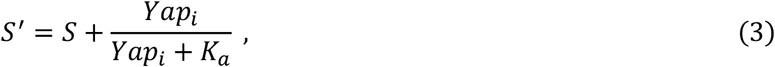

where the variable *Yap*_*i*_ represents an expression level of Yap. Furthermore, the parameter *S* represents a basal transcription rate of Cdx2, and the parameter *K*_*a*_ represents the Half-maximal concentration of Yap for Cdx2 induction. The model of Yap is described by Equation (4). This equation expresses the concentration dynamics of Yap [Caluwé et al., 2019]. Because Yap is phosphorylated by phosphorylated Lats, they cannot enter the nucleus:

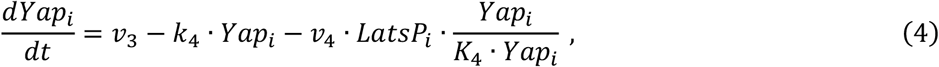

where the parameter *v*_3_ reflects a production rate of Yap, the parameter *k*_4_ is the kinetic constant of basal degradation of Yap, and the parameter *v*_4_ is a phosphorylation rate of Yap. Furthermore, the variable *LatsP*_*i*_ represents an expression level of phosphorylated Lats. Eq. (4) consists of three terms: the first term represents the production level of Yap, the second term represents the degradation of Yap, and the third term represents the phosphorylation of Yap.

Phosphorylated Lats is promoted by the Nf2-Amot complex. The model of phosphorylated Lats is described by Equation (5), where the parameter *v*_1_ reflects a phosphorylation rate of Lats, the parameter *K*_1_ is the Michaelis-Menten constant for Lats phosphorylation, the parameter *v*_2_ is a dephosphorylation rate of phosphorylated Lats, and the parameter *K*_2_ is the Michaelis-Menten constant for of phosphorylated Lats dephosphorylation. The Michaelis-Menten constant is a coefficient which used as a guideline for enzyme reactions. The smaller value shows the higher affinity between the enzyme and the substrate. Eq. (5) expresses the concentration dynamics of phosphorylated Lats:

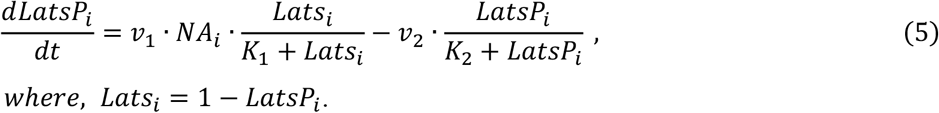

In Eq. (5), the variable *NA*_*i*_ represents an expression level of the Nf2-Amot complex. Furthermore, the variable *Lats*_*i*_ represents an expression level of Lats. Eq. (5) consists of two terms: the first term represents the phosphorylation of Lats by the Nf2-Amot complex, and the second term represents the dephosphorylation of phosphorylated Lats.

Nf2 forms the Nf2-Amot complex on E-cadherin and does not form the complex on Par-aPKC. We focus on the Nf2-Amot complex since Nf2 forms the complex and afterward acts on the phosphorylation cascade. Equation (6) expresses the concentration dynamics of the Nf2-Amot complex. The proposed model referred to the dynamics of the previous model to build the dynamics. The model of the Nf2-Amot complex is described by Eq. (6), where the parameter *v*_5_ reflects a production rate of the Nf2-Amot complex, the parameter *k*_5_ is the kinetic constant of basal degradation of the Nf2-Amot complex, the parameter *v*_6_ is a degradation rate of the Nf2-Amot complex, and the parameter *K*_5_ is the Michaelis-Menten constant for the Nf2-Amot complex degradation:

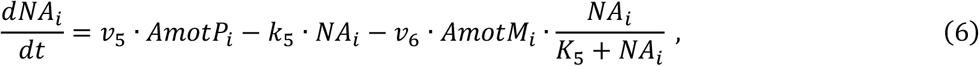

where the variable *AmotP*_*i*_ represents an expression level of Amot is phosphorylated, and the variable *AmotM*_*i*_ represents an expression level of Amot is trapped by the Par-aPKC. Eq. (6) consists of three terms: the first term represents the production of the Nf2-Amot complex, the second term represents the degradation of the Nf2-Amot complex, and the third term represents the formation inhibition of the Nf2-Amot complex.

Amot is distributed on E-cadherin, but when Par-aPKC is present, Amot is sequestered from E-cadherin and is trapped by Par-aPKC. While Equation (7) expresses the concentration dynamics of phosphorylated Amot, Equation (8) expresses the concentration dynamics of Amot trapped by Par-aPKC [Caluwé et al., 2019]. The model of AmotP is described by Eq. (7), where the parameter *k*_*a*1_ reflects the Amot/E-cadherin association kinetic constant, and the parameter *k*_*d*1_ is the Amot/E-cadherin dissociation kinetic constant. The model of AmotM is described by Eq. (8), where the parameter *k*_*a*2_ reflects the Amot/Par-aPKC association kinetic constant, and the parameter *k*_*d*2_ is the Amot/Par-aPKC dissociation kinetic constant. The differential equations are described as:

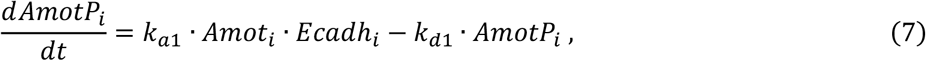

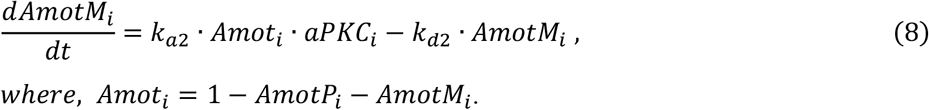

In these equations, the variable *Amot*_*i*_ represents an expression level of Amot, the variable *Ecadh*_*i*_ represents an expression level of E-cadherin, and the variable *aPKC*_*i*_ represents an expression level of Par-aPKC. Eq. (7) consists of two terms: the first term represents Amot/E-cadherin association, and the second term represents Amot/E-cadherin dissociation. Eq. (8) also consists of two terms: the first term represents Amot/Par-aPKC association, and the second term represents Amot/Par-aPKC dissociation.

In the embryo, cells recognize their position through cell-cell contacts. Cell-cell contacts are represented by the concentration dynamics of Par-aPKC/E-cadherin. While Equation (9) expresses the concentration dynamics of Par-aPKC, Equation (10) expresses the concentration dynamics of E-cadherin. These proposed models use the dynamics of the previous model as references. These models are described using the following coefficients: the parameter *W* reflects the basal transcription rate of Par-aPKC (outer cell) or E-cadherin (inner cell), the parameter *X* is the basal transcription rate of E-cadherin (outer cell) or Par-aPKC (inner cell), the parameter *Y* is Par-aPKC/E-cadherin reciprocal inhibition strength, the parameter *z* is a transcription rate, the parameter *Z* is threshold constant of auto-activation and cross-inhibition, and the parameter *Q* is gene expression timing of Par-aPKC/E-cadherin. Concerning the outer cell, Eq. (9) uses the parameter W, and Eq. (10) uses the parameter X. On the other hand, concerning the inner cell, Eq. (9) uses the parameter X, and Eq. (10) uses the parameter W. By changing the value of the parameter *Q* in these differential equations, it was possible to simulate the diverse concentrations dynamics of Par-aPKC and E-cadherin.

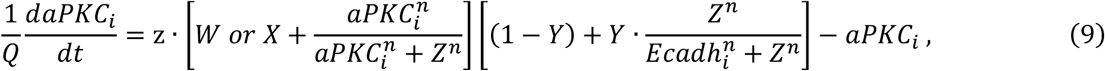

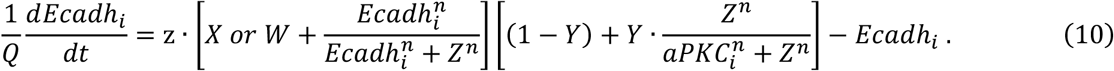

Eq. (9) and Eq. (10) consist of two terms: the first bracket in the first term represents self-amplification, the second bracket in the first term represents mutual inhibition, and the second term represents Par-aPKC/E-cadherin degradation, respectively.

Using the above Equations (1) to (10), we conduct two experiments to show that the proposed model outperforms the previous one and guide us to our assumption, i.e., the Nf2-Amot complex regulates the cell differentiation of TE and ICM. The next subsection determines the parameters *v*_5_, *k*_5_, *v*_6_, and *K*_5_ in the model of the Nf2-Amot complex.

### The parameters selection in the model of the Nf2-Amot complex

The parameters selection referred to the ranges of the parameters in the previous model. The parameter *v*_5_ reflects a production rate of the Nf2-Amot complex, the parameter *k*_5_ is the kinetic constant of basal degradation of the Nf2-Amot complex, the parameter *v*_6_ is a degradation rate of the Nf2-Amot complex, and the parameter *K*_5_ is the Michaelis-Menten constant for the Nf2-Amot complex degradation. To determine these parameters, we referred to the parameter in the previous model which represents a production rate, the kinetic constant of basal degradation, a degradation rate, and the Michaelis-Menten constant, respectively. Thus, the range of each parameter is as follows:

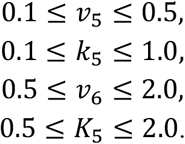

Within these ranges, the experiment determined an ideal combination of the parameters. The ideal combination of the parameters is a combination in which position-specific cell differentiation is expressed. The experiment used the concentration dynamics of Par-aPKC/E-cadherin shown in **Fig.3** and searched for the ideal combination of parameters. During the searching process of the parameters, we changed the value of each parameter in the Nf2-Amot complex, i.e., the Eq. (6), by 0.1, and the results expressed three-dimensional parameter spaces shown in **Fig.4**. In **Fig.4**, the blue dots show the combinations of the parameters that predominantly expressed Cdx2 in the outer cell. The red dots show the combinations of the parameters which predominantly expressed Oct4 in the inner cell. On the other hand, the green dots show the ideal combinations of the parameters. The result shows that the green dots increase with the increase of parameter *v*_5_. Additionally, *K*_5_ reduced the level of the Nf2-Amot complex more strongly than *v*_6_, as expected from the model.

**Fig. 3.**
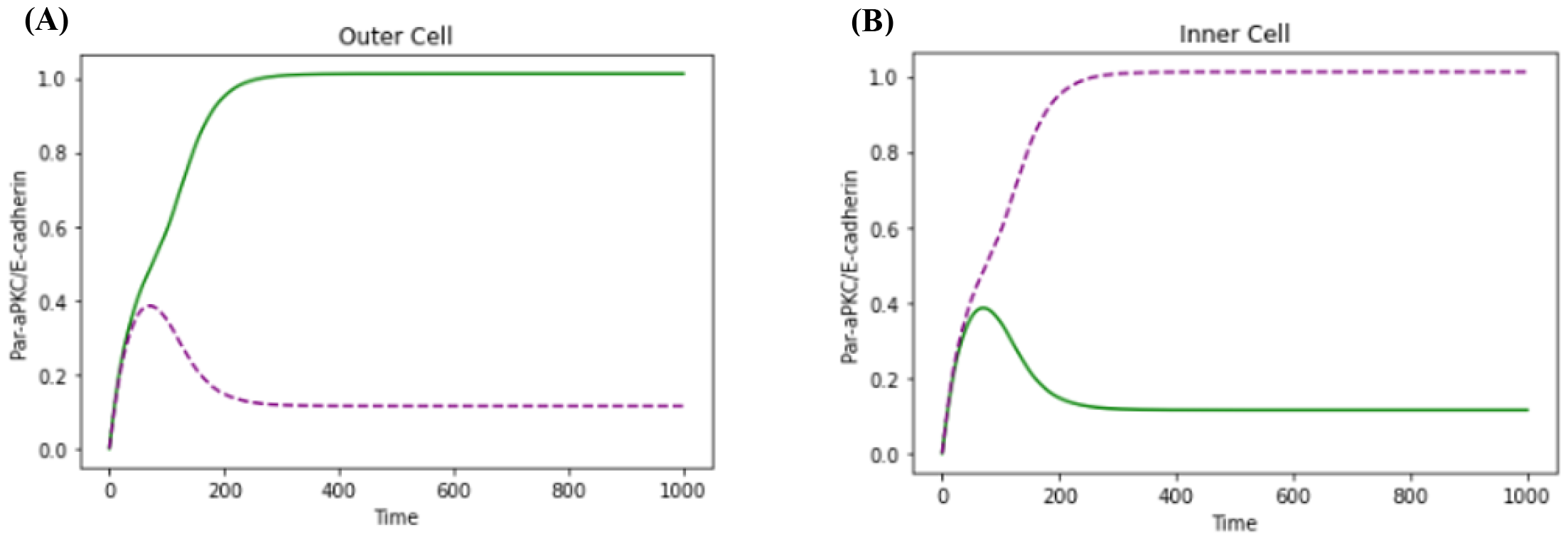
Concentration dynamics of mutually inhibitory Par-aPKC/E-cadherin [Caluwé et al., 2019]. These temporal evolutions of Par-aPKC/E-cadherin are initially upregulated during compaction. In Equations (9) and (10), the value of a parameter Q is 0.03. The green line represents a concentration change of Par-aPKC. The purple line represents a concentration change of E-cadherin. (A) Par-aPKC upregulates and E-cadherin slightly downregulates in the outer daughter cell. (B) Par-aPKC downregulates and E-Cadherin upregulates in the inner daughter cell.

**Fig. 4.**
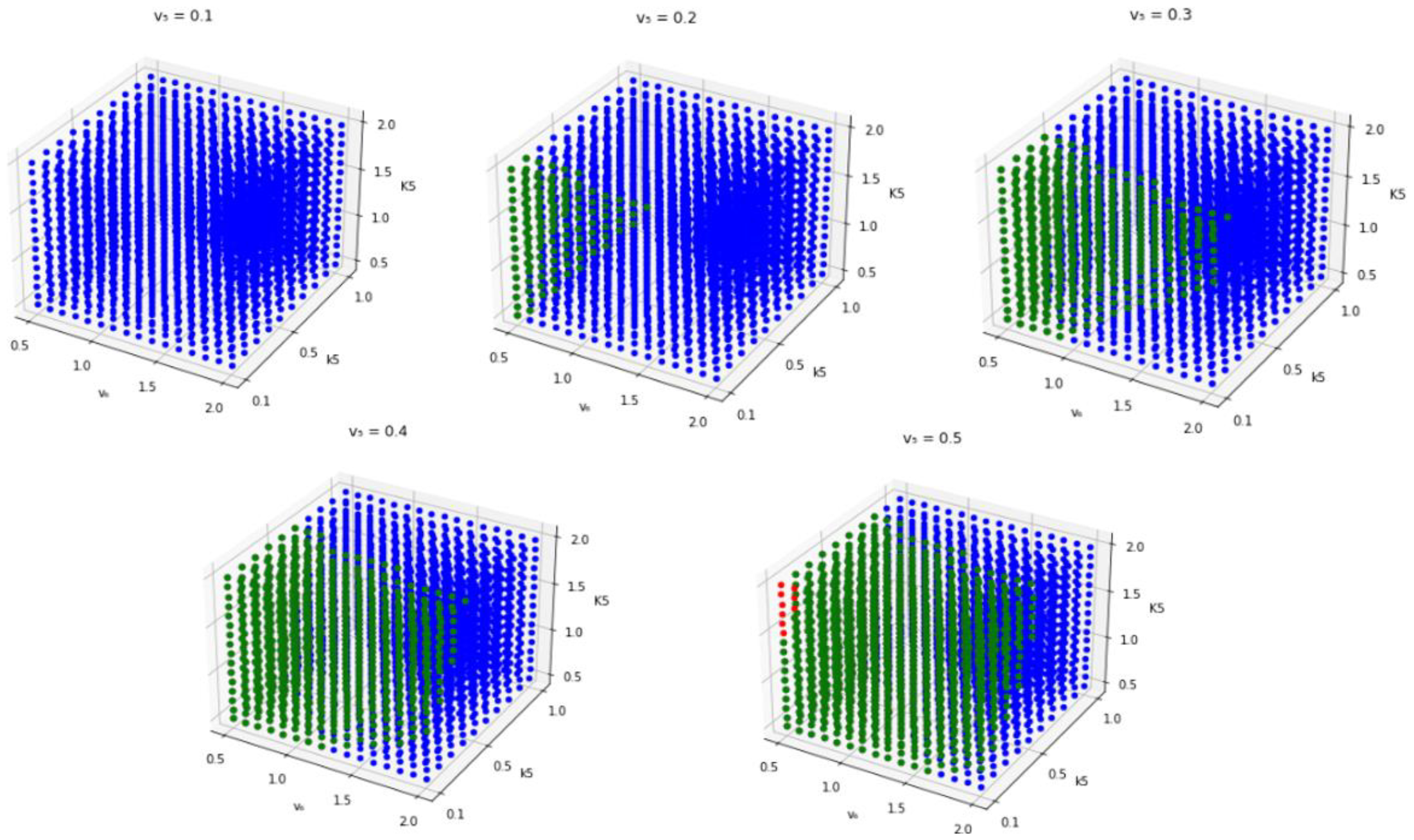
Parameter decision in the differential equation of the Nf2-Amot complex. We searched for combinations of four parameters (v_5_, k_5_, v_6_, K_5_) that enable Cdx2/Oct4 expression depending on the location in embryos. The blue dots indicate the combinations of the parameters that enabled Cdx2 to be predominantly expressed in the dynamics of Par-aPKC/E-cadherin, assuming the outer cells in embryos. The red dots indicate the combinations of the parameters that enabled Oct4 to be predominantly expressed in the dynamics of Par-aPKC/E-cadherin, assuming the inner cells in the embryo. The green dots indicate the combination of parameters that allowed the predominant expression of ideal transcription factors, i.e., Cdx2 and Oct4, in the outer and inner cells of embryos. If not reproducing the expression of transcription factors, i.e., Cdx2 and Oct4, according to its position, the point corresponding to the parameter combination is missing.

The parameters of the model are determined from among ideal combinations of the parameters. Any green dot at least expressed the position-specific cell differentiation in the cell-cell contacts shown in **Fig.3**. The paper used the following combination randomly selected from among the green dots:

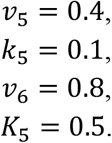

### High Cdx2 expression in dominant-negative form of the Nf2-Amot complex

We experimented with concentration restrictions on the Nf2-Amot complex to confirm whether the dynamics of the proposed model follow that of the embryo *in vivo*. To assess the dynamics in the proposed model, the experiment compared simulation with the biological observation. The observation is that the dominant-negative form of Nf2 (dnNf2) causes Cdx2 misexpression in the inner cell [Cockburn et al., 2013]. Here, the model confirmed whether inhibition of the Nf2-Amot complex causes high Cdx2 expression in the inner cell.

A low value of variable *NA*_*i*_ *in silico* expressed the inhibition of the Nf2-Amot complex. The experiment used the value of the Nf2-Amot complex by changing the variable to constant to clarify the difference in the dynamics of each value. The value was investigated separately for (i) 0 and (ii) the range of non-zero values: (i) **Fig.5A** and **5B** show the result when the value is 0; (ii) **Fig.5C** and **5D** show the result when the value is 0.33, which is the maximum value when high Cdx2 expression in the inner cell. The experiment used the concentration dynamics of Par-aPKC/E-cadherin of 100 patterns. The next subsection described the diverse dynamics of Par-aPKC/E-cadherin.

**Fig. 5.**
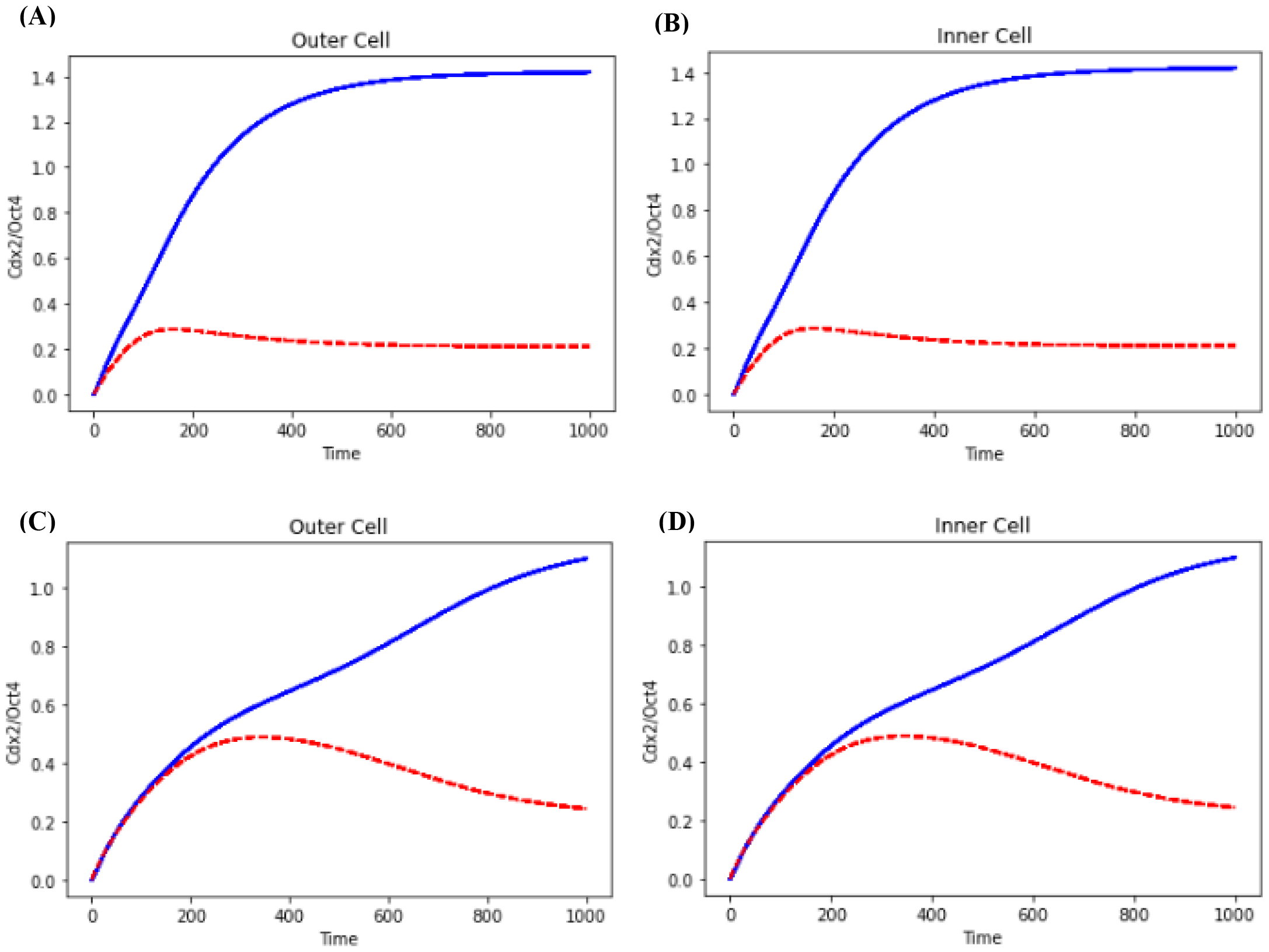
Temporal evolution of Cdx2 and Oct4 in dnNf2 with a computational model. These temporal evolutions represent the concentration dynamics of Cdx2 and Oct4. While the blue line represents the concentration level of Cdx2, the red line represents the concentration level of Oct4. (A, B) *NA*_*i*_ = 0. (C,D) *NA*_*i*_ = 0.33.

The formation inhibition of the Nf2-Amot complex promoted the high expression level of Cdx2 *in silico*. When (i) the value is 0, the components of the Hippo pathway did not phosphorylate, and then the inner cell did not differentiate ICM (**Fig.5B**). Furthermore, while the cell-cell contacts express the diverse concentration dynamics of Par-aPKC/E-cadherin, the results show the same concentration dynamics of Cdx2/Oct4 (**Fig.5A** and **5B**).

The experiment specified the range of *NA*_*i*_ which is able to express dnNf2. When (ii) the range of non-zero values, i.e., the range of values from 0.01 to 0.33 (interval: 0.01), the model was able to reproduce the observation. Furthermore, the same as the results of **Fig.5A** and **5B, Fig.5C** and **5D** show the same concentration dynamics of Cdx2/Oct4. From (i) and (ii), when 0 ≤ *NA*_*i*_ ≤ 0.33, the proposed model was able to simulate that dnNf2 causes Cdx2 misexpression in the inner cell. The proposed model, thus, follows the dynamics of the embryo *in vivo*.

### Concentration dynamics of Cdx2/Oct4 by the existence of the Nf2-Amot complex

The experiment examined how the existence of the Nf2-Amot complex influences the expression of Cdx2 and Oct4. The comparative experiment intends to infer the role of the Nf2-Amot complex. We compared the proposed model and the previous model using the diverse concentration dynamics of Par-aPKC/E-cadherin (Eq. (9) and Eq. (10)). By changing parameter *Q*, i.e., the gene expression timing of Par-aPKC/E-cadherin, the experiment expressed the diverse concentration dynamics. We determined the range of *Q* since the difference between each dynamics becomes less likely to change the results as it increases by a preliminary experiment. Then, the range of *Q* was as:

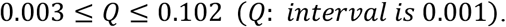

The range expresses the concentration dynamics of 100 patterns (**Fig.6A** and **6B**). Using these dynamics, we conducted a comparative experiment to observe the robustness.

**Fig. 6.**
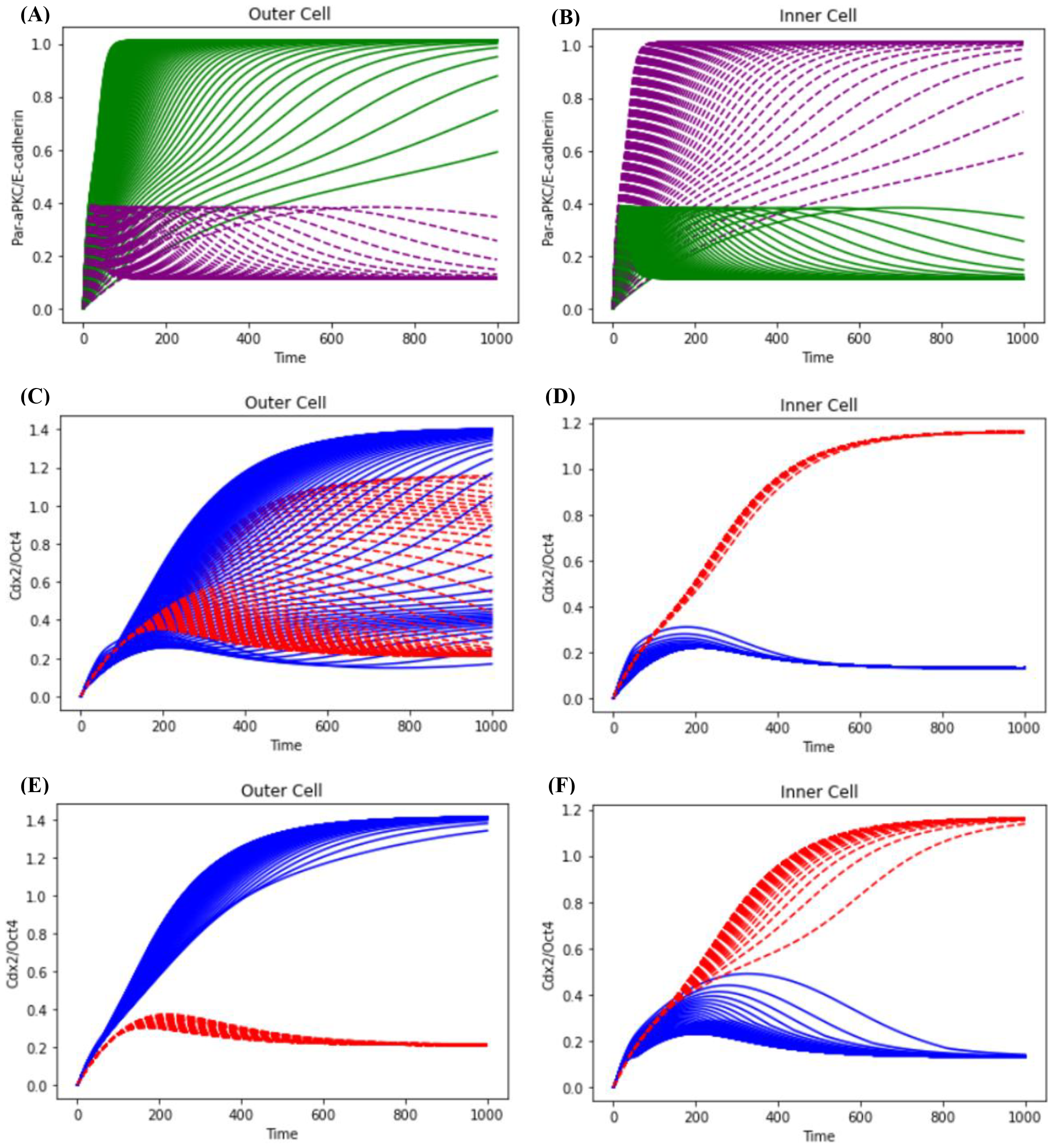
Temporal evolution of Cdx2/Oct4 by the concentration dynamics of Par-aPKC/E-cadherin. (A, B) Temporal evolution of Par-aPKC and E-cadherin. These express the cell-cell contacts. The green line represents the concentration dynamics of Par-aPKC. The purple line represents the concentration dynamics of E-cadherin. (C-F) The blue line represents the concentration dynamics of Cdx2. The red line represents the concentration dynamics of Oct4. (C, D) The previous model results in temporal evolutions of Cdx2 and Oct4. (E, F) The proposed model results in temporal evolutions of Cdx2 and Oct4.

The concentration dynamics of Par-aPKC/E-cadherin located in the cell in the embryo. These models, Eq. (9) and Eq. (10), express the dynamics in the daughter cell. Thus, the dynamics initially increase both Par-aPKC/E-cadherin during compaction. After cell division, each cell increases and decreases Par-aPKC/E-cadherin depending on their location. While the cell increases Par-aPKC and decreases E-cadherin in the outer cell (**Fig.6A**), the cell increases E-cadherin and decreases Par-aPKC in the inner cell (**Fig.6B**). Hereinafter, the paper refers to the outer daughter cell and the inner daughter cell as the outer cell and the inner cell, respectively.

The paper used the previous model as a model, in which the Nf2-Amot complex does not exist but regulates Cdx2/Oct4. It’s not that the previous model inhibits the Nf2-Amot complex. Namely, the phosphorylation cascade is regulated without considering the Nf2-Amot complex (**Fig.2A**). The model, thus, can express in specific cell-cell contact conditions. Comparing the proposed model and the previous model, the difference between the two implies the role of the Nf2-Amot complex.

The Nf2-Amot complex strengthened the expression of Cdx2 in the outer cell. The results show the concentration dynamics of Cdx2/Oct4 in the outer cell and the inner cell (**Fig.6C-6F**). The previous model showed the concentration dynamics of Cdx2/Oct4 (**Fig.6C** and **6D**). The outer cell observed that the concentration dynamics of Oct4 are higher than that of Cdx2 (**Fig.6C**). On the other hand, the inner cell observed that the concentration dynamics of Oct4 are higher than that of Cdx2, as the dynamics of the embryo *in vivo* (**Fig.6D**). The proposed model showed the concentration dynamics of Cdx2/Oct4 (**Fig.6E** and **6F**). While the outer cell observed that the concentration dynamics of Cdx2 are higher than that of Oct4 (**Fig.6E**), the inner cell observed that the concentration dynamics of Oct4 are higher than that of Cdx2 (**Fig.6F**), as the dynamics of the embryo *in vivo*.

As the dominant timing for Par-aPKC or E-cadherin gets late, the previous model promoted the Oct4 expression in the outer cell. To make a detailed observation of the dynamics shown in **Fig.6**, we extracted the Par-aPKC/E-cadherin dynamics of four patterns and then conducted the comparative experiment. The four dynamics were generated by setting the values of *Q* to 0.003, 0.007, 0.020, and 0.102. These values were determined in the range of *Q*, i.e., minimum, maximum, and random sampling of the dynamics which largely differ from the dynamics of minimum and maximum. As is obvious from **Fig.6**, once again, both models observed that the concentration dynamics of Oct4 are higher than that of Cdx2 in the inner cell (**Fig.7**). On the other hand, as the timing of the stable steady states of Par-aPKC/E-cadherin gets late, the previous model was not able to reproduce position-specific cell differentiation in the outer cell (**Fig.7A, 7B**, and **7C**).

**Fig. 7.**
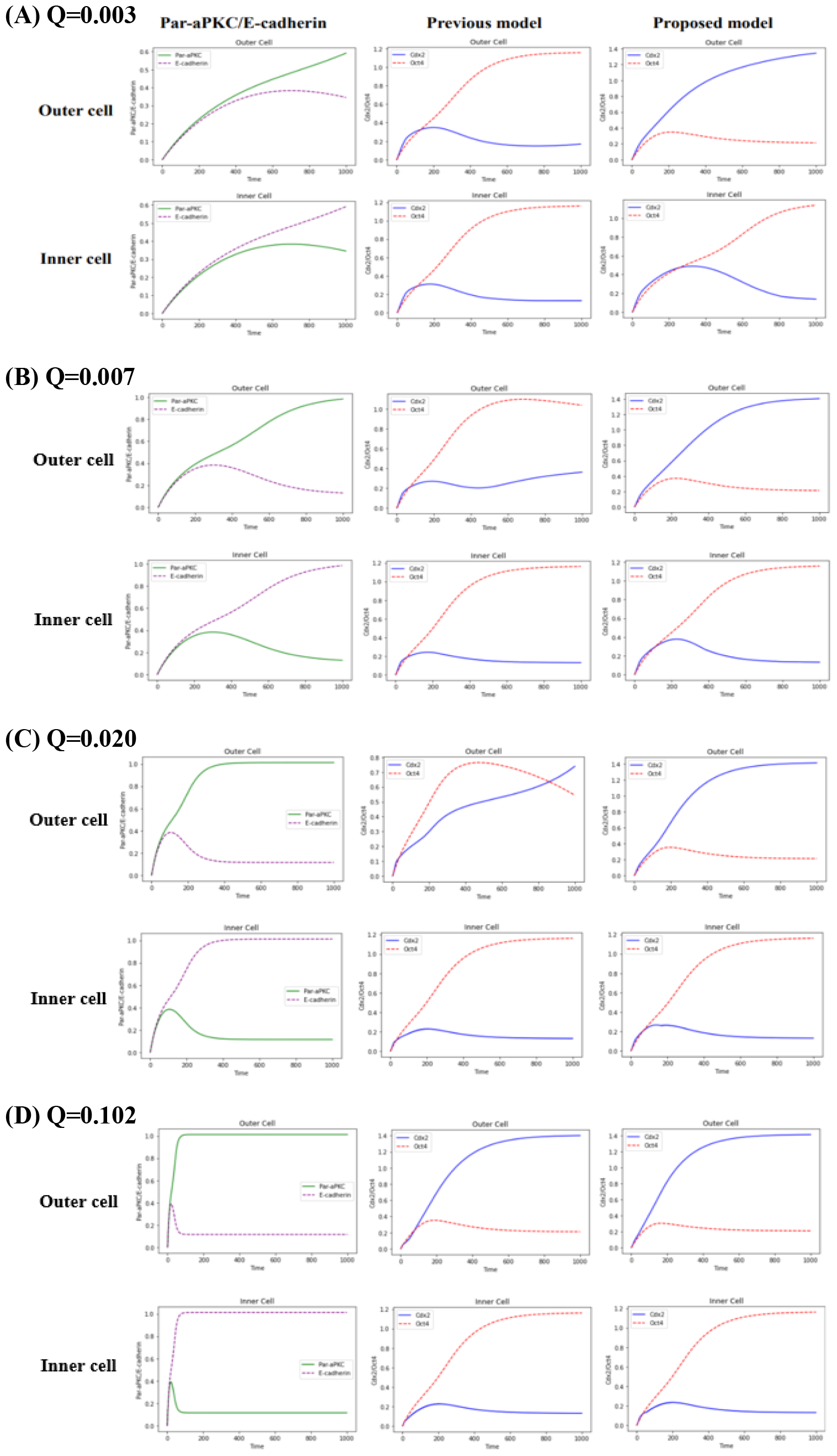
Temporal evolution of Cdx2/Oct4 by Par-aPKC/E-cadherin dynamics of 4 patterns. In the range of Q, these values were determined by minimum, maximum, and random sampling of the dynamics which largely differ from the dynamics of minimum and maximum. The green line represents the concentration dynamics of Par-aPKC. The purple line represents the concentration dynamics of E-cadherin. The blue line represents the concentration dynamics of Cdx2. The red line represents the concentration dynamics of Oct4. (A) Q = 0.003. (B) Q = 0.007. (C) Q = 0.020. (D) Q = 0.102.

## DISCUSSION

In developmental biology, it is important to understand the regulation mechanism of the embryo. The early mammalian embryo has long been investigated in developmental biology. In this development, concurrently with compaction cells acquire an apical-basal polarity [Toyooka et al., 2016; Cockburn et al., 2010], and cells adapt to the cell-cell contact conditions using the Hippo signaling pathway. However, this regulation mechanism is still due to controversy and the details have not been clarified. The paper assumes that the Nf2-Amot complex plays the regulatory mechanism of the Hippo signaling pathway. To confirm the assumption, this paper discusses three main things. The first section discusses the reliability of a computational model. This confirms whether the computational model can phenomenologically reproduce biological knowledge. The second section focuses on the interactions between components in the Hippo signaling pathway with the results (**Fig.6** and **Fig.7**). The interactions among the components suggest that the presence of the Nf2-Amot complex is important for proper the cell differentiation. The third section discusses presence of the Nf2-Amot complex influences the cell differentiation of TE and ICM. Here, the models with and without the Nf2-Amot complex were compared to reveal the functions of the Nf2-Amot complex. The absence of the Nf2-Amot complex causes over-phosphorylation resulting in misexpression of Oct4 in the outer cell. This result suggests that the Nf2-Amot complex regulates the Hippo signaling pathway. Finally, we discuss that the Nf2-Amot complex regulates the phosphorylation cascade of the Hippo signaling pathway to make the cell differentiation properly in early mammalian development, i.e., maintains the robustness in the embryo.

### Reliability of a computational model

The proposed model phenomenologically reproduces the current biological knowledge regarding the early mammalian embryo as discussed below. First, the living cells of the embryo produce Cdx2 and Oct4 by following the cell-cell contact conditions, i.e., the concentration dynamics of Par-aPKC and E-cadherin. On the other hand, the proposed model reproduces the expression of Cdx2 and Oct4 following the cell locations in the embryo (**Fig.6** and **Fig.7**). Second, while dnNf2 causes Cdx2 misexpression in the inner cell *in vivo* [Cockburn et al., 2013], the model reproduces this biological knowledge (**Fig.5**). Thus, it can say that the proposed model phenomenologically best reproduces the living cells of the early mammalian embryo. From these facts, we can infer that the cell behavior *in vivo* can be reproduced by the proposed model of the cells *in silico*. The next subsection discusses the results shown in **Fig.6** and **Fig.7** to understand why the results were different.

### Regulation of the Hippo signaling pathway and improvement of the robustness by the Nf2-Amot complex

The results of experiments shown in **Fig.6** and **Fig.7** express that the Nf2-Amot complex improves the robustness by regulating the phosphorylation cascade of the Hippo signaling pathway. The paper focuses on the robustness of computational models since the living cells in the embryo have extraordinary adaptability and robustness [Holmes et al., 2017; Toyooka, 2020]. The robustness of the model can be observed as the predominance of Cdx2 or Oct4 depends on cellular locations, i.e., diverse cell-cell contact surfaces, within the embryo. The comparative experiment demonstrated that the Nf2-Amot complex regulates the expression of Cdx2 and Oct4 in diverse cell-cell contact conditions and improves robustness of the system (**Fig.6** and **Fig.7**). To examine the mechanism of how the Nf2-Amot complex improves robustness, we analyzed the results of **Fig.6** and **Fig.7** in detail. To investigate how the Nf2-Amot complex regulates the expression of Cdx2 and Oct4, we confirmed the model structure shown in **Fig.2**.

Incorporating the Nf2-Amot complex into the model allows cell differentiation depending on diverse cell-cell contacts. The previous model has a structure that *AmotP*_*i*_ directly phosphorylates Lats, and *AmotM*_*i*_ inhibits *AmotP*_*i*_. Namely, the previous model is less likely to influence the production of *LatsP*_*i*_ by *AmotM*_*i*_ (**Fig.2A**). This means that the structure of the model itself favors phosphorylation results in leverage to the inner cell. On the other hand, the proposed model is a structure that the Nf2-Amot complex directly phosphorylates Lats, and it is simultaneously influenced by *AmotP*_*i*_ and *AmotM*_*i*_ (**Fig.2B**). This suppresses phosphorylation of Lats by *AmotM*_*i*_ and reproduces the cell differentiation in the outer cell. Furthermore, the proposed model can reproduce dominant expressions properly not only in the outer cell but also in the inner cell (**Fig.6E** and **6F**). With these facts, the result suggests that the Nf2-Amot complex plays a role in reflecting the concentration dynamics of Par-aPKC and E-cadherin to regulate the Hippo signaling pathway. The next subsection focuses on the amount of *LatsP*_*i*_ to examine the existence of the Nf2-Amot regulates the phosphorylation cascade of the Hippo signaling pathway. An additional experiment conducted in the next subsection is difficult to examine in the living cells using the model.

### A regulation function of phosphorylation in early mammalian development

The Nf2-Amot complex regulates the maximum level of phosphorylated Lats (**Fig.8**). In the experiments, the proposed model could reproduce differentiations of both cells while the previous model could reproduce only the inner cell. Thereby, we thought there were clues in the outer cell. We conducted an additional experiment to confirm the concentration dynamics of phosphorylated Lats (*LatsP*_*i*_) in the outer cell. The additional experiment used the concentration dynamics of 100 patterns (**Fig.6A** and **6B**) crested for assuming diverse cell-cell contact surfaces. The result reveals that the maximum concentration of phosphorylated Lats in the proposed model is lower than that in the previous model (**Fig.8**). This means that the existence of the Nf2-Amot complex prevents the excessive phosphorylation of Lats.

**Fig. 8.**
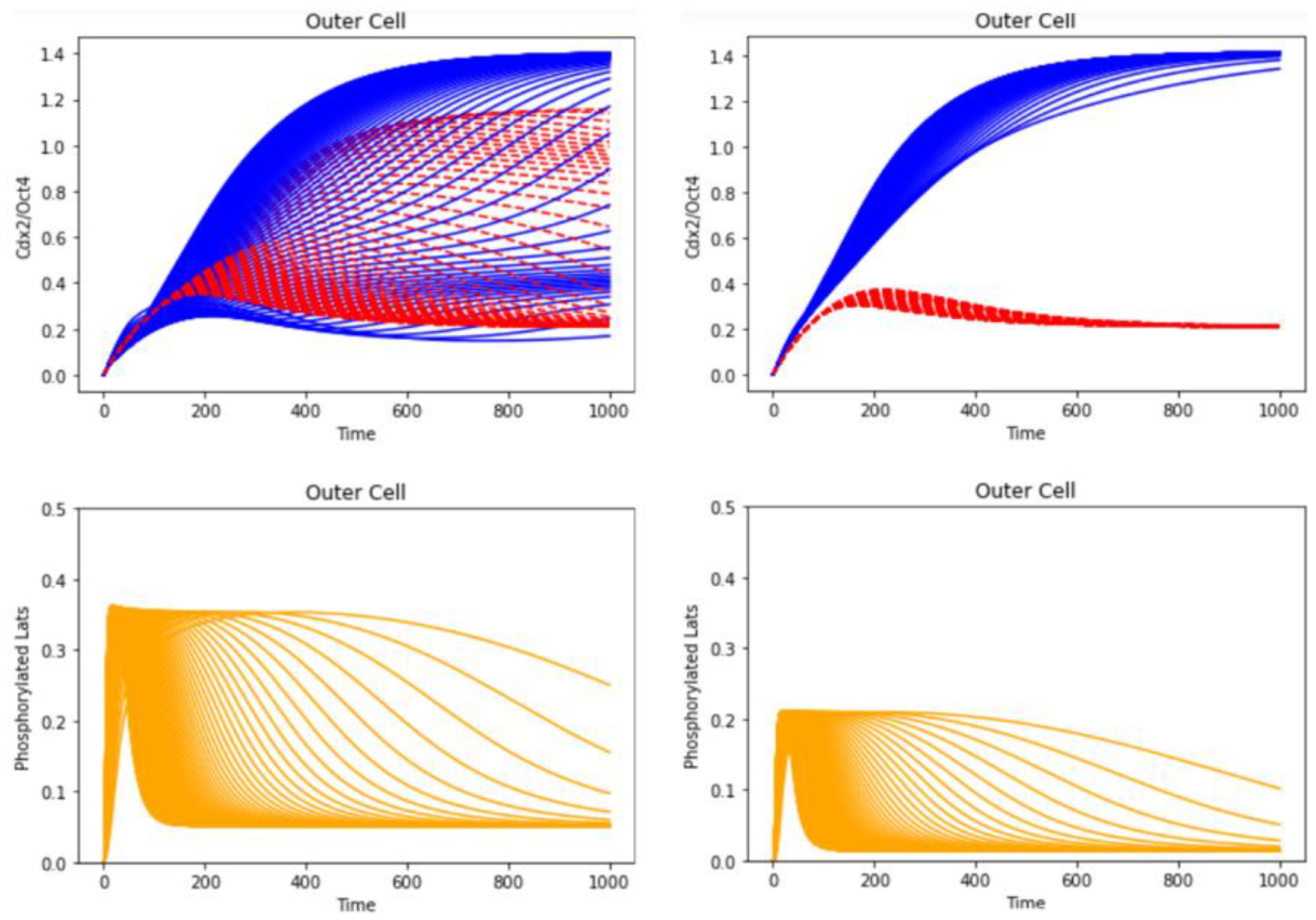
The concentration dynamics of Cdx2/Oct4 and phosphorylated Lats in the outer cell. The blue line represents the concentration dynamics of Cdx2, the red line is the concentration dynamics of Oct4, and the orange line is the concentration dynamics of phosphorylated Lats. The previous model shows the concentration dynamics of Cdx2/Oct4 (upper left) and phosphorylated Lats (bottom left). The proposed model shows the concentration dynamics of Cdx2/Oct4 (upper right) and phosphorylated Lats (bottom right).

We suggest that the Nf2-Amot complex improves the robustness of the whole embryo by regulating the phosphorylation cascade. On the outer cell, the paper demonstrated that the Nf2-Amot complex regulates the cell differentiation of TE *in silico* (**Fig.6, Fig.7**, and **Fig.8**). Furthermore, on the inner cell, the Nf2-Amot complex regulates the cell differentiation of ICM from the experiment (**Fig.5** and **Fig.6**) and the biological knowledge [Cockburn et al., 2013]. From these facts, the paper derives the assumption that the Nf2-Amot complex regulates the cell differentiation of TE and ICM.

### Limitations and prospect of the computational model

Results obtained from the computational model are only inferences. The computational model expresses the behavior of intracellular substances in the embryo, and each differential equation expresses the dynamics of a target protein. Namely, previous studies on theoretical biology have modeled based on observations, e.g., fluorescence observations. Thus, Theoretical approaches can simulate a target mechanism, and infer various properties and roles of proteins. However, the result is only an inference since it is difficult to reproduce all components, all mechanisms (chemical reactions), and detailed expression timing in a cell.

Theoretical approaches provide novel assumptions for biology. Computational models contribute understanding of intracellular mechanisms from dynamical and phenomenological perspectives. This is because the computational model is simplification of the target system and expresses its essence. The model preserves the properties of the target part of the real system, so it is possible to understand the system or to infer the behavior of the real system. Thus, insights from computational models provide useful assumptions for biology.

The proposed model is improved by the selection of the parameters based on biological observations, and thereby the role of the Nf2-Amot complex can be more accurately inferred. The proposed model is an improvement on the previous model from the perspective of improving robustness, and the correctness of the model is demonstrated by reproducing the observation of the previous study [Caluwé et al., 2019]. However, the parameters selection of the proposed model is determined by the combination of parameters in which the position-specific cell differentiation is expressed (**Fig.4**), so the model does not consider the concentration dynamics of the Nf2-Amot complex (Nf2) *in vivo*. Therefore, future work is going to improve the model by the parameters selection based on biological observations.

## CONCLUSION

In conclusion, the Nf2-Amot complex is thought to lead cell differentiation properly in early mammalian development by dynamically regulating the phosphorylation in both the inner cell and the outer cell. the paper quantifies the concentration dynamics of the main components in cell differentiation and the Nf2-Amot complex effects on early mammalian development to infer the function of the Nf2-Amot complex. This approach focused on the Nf2-Amot complex since the cell differentiation of the inner cell needs to regulate the Hippo signaling pathway from the biological knowledge in the early mammalian development. We proposed a novel computational model by constructing the model of the Nf2-Amot complex and integrating it into the previous model. Consistent with the biological knowledge, inhibition of the Nf2-Amot complex inhibited the differentiation of ICM in the proposed model. Furthermore, comparing the previous model and the proposed model in diverse cell-cell contact surfaces, the result showed that the proposed model had improved robustness. Thus, based on these results and the biological knowledge, the paper provides the hypothesis that the Nf2-Amot complex regulates the phosphorylation cascade of the Hippo signaling pathway to make cell differentiation properly in early mammalian development.

